# Autophagy in T cells from aged donors is maintained by spermidine, and correlates with function and vaccine responses

**DOI:** 10.1101/2020.06.01.127514

**Authors:** Ghada Alsaleh, Isabel Panse, Leo Swadling, Hanlin Zhang, Alain Meyer, Janet Lord, Eleanor Barnes, Paul Klenerman, Christopher Green, Anna Katharina Simon

## Abstract

Older adults are at high risk for infectious diseases such as the recent COVID-19 and vaccination seems to be the only long-term solution to the pandemic. While most vaccines are less efficacious in older adults, little is known about the molecular mechanisms that underpin this. Autophagy, a major degradation pathway and one of the few processes known to prevent aging, is critical for the maintenance of immune memory in mice. Here, we show induction of autophagy is specifically induced in human vaccine-induced antigen-specific T cells *in vivo*. Reduced IFNγ secretion by vaccine-induced T cells in older vaccinees correlates with low autophagy. We demonstrate in human cohorts that levels of the endogenous autophagy-inducing metabolite spermidine, fall with age and supplementing it *in vitro* recovers autophagy and T cell function. Finally, our data show that endogenous spermidine maintains autophagy via the translation factor eIF5A and transcription factor TFEB. With these findings we have uncovered novel targets and biomarkers for the development of anti-aging drugs for human T cells, providing evidence for the use of spermidine in improving vaccine immunogenicity in the aged human population.

## Introduction

The outbreak of coronavirus disease 2019 (COVID-19) caused a great threat to world-wide public health in 2020 with the majority of deaths occurring in older adults. The development of effective treatments and vaccines against COVID-19 is now more than ever becoming a pressing and urgent challenge to overcome^1,2^. However the successful vaccination of the elderly against pathogens is considered one of the big challenges in our society^3,4^. Immunosenescence, which is characterized by poor induction and recall of B and T memory responses upon exposure to new antigens, can lead to reduced immune responses following immunization of older adults. While most vaccines are less immunogenic and effective in the older population ^3^, little is known about the molecular mechanisms that underpin immune senescence. Autophagy is thought to be one of the few cellular processes that underlie many facets of cellular ageing including immune senescence ^5^. By delivering unwanted cytoplasmic material to the lysosomes for degradation, autophagy limits mitochondrial dysfunction and accumulation of reactive oxygen species (ROS) ^6^. Autophagy degrades protein aggregates that accumulate with age and its age-related decline could contribute to “inflammaging” ^7^, the age-related increase in inflammatory cytokines in in blood and tissue. Loss of autophagy strongly promotes production of the inflammatory cytokines TNFa, IL-6 and IL1-β ^8,9^. We previously found autophagy levels decline with age in human peripheral CD8^+^ T cells ^10^. Deletion of key autophagy genes leads to a prematurely aged immune phenotype, with loss of function in mouse memory CD8^+^ T cells ^11 12^, hematopoietic stem cells ^13^, and macrophages ^9^ with a myeloid bias ^13^. In addition, we find in autophagy-deficient immune cells the same cellular phenotype that cells display in older organisms; they accumulate ROS and damaged mitochondria ^9,12^.

Importantly, we can improve CD8^+^ T memory responses from aged mice with spermidine ^12^, an endogenous metabolite synthesized from arginine. It was shown in yeast and other model organisms that spermidine extends life-span via increased autophagy ^14^. Several downstream mechanisms of spermidine-induced autophagy have been described in mice, including the inhibition of histone deacetylases ^14^. Recently we uncovered a novel pathway in which spermidine lends a residue for the hypusination of the translation factor eIF5a, which is necessary for the translation of a three proline motif present in the master transcription factor of autophagy and lysosomal biogenesis, called TFEB ^15^. We demonstrated this pathway operates in activated B cells, which upon activation have an unusually high protein synthesis rate, owing to the high production of immunoglobulins. It is likely that B cells may be particularly reliant on the unfolded protein response, the proteasome, and autophagy, to cope with this high rate of protein synthesis. B cells may have evolved special coping strategies including the translational signalling for autophagy via eIF5A and TFEB. We therefore sought to extend our findings to another immune cell type, CD8+ T cells, to investigate whether this pathway may be conserved in a related adaptive immune subset and possibly broadly applicable.

Here we show for the first time that autophagy is indeed highly active in human CD8+ T cells after the *in vivo* encounter of antigens in donors from two different experimental vaccination trials. Our data show that polyamine levels fall with age in peripheral mononuclear cells. When supplemented with spermidine, the dysfunctional autophagic flux can be rejuvenated in CD8+ T cells from old donors, and levels of the important effector molecules IFNγ and perforin are enhanced as a consequence. Moreover, autophagy and effector function are maintained by spermidine in T cells from young donors. Lastly, in human CD8+ T cells we show that spermidine signals via eIF5A and TFEB to maintain autophagy levels. This study demonstrates that the function of human CD8+ T cells can be improved with spermidine. Taken together with our previous work on B cells, this leads us to the hypothesis that both T and B cell responses to infections and vaccinations are exquisitely reliant on sufficient autophagy levels, which are maintained by intracellular spermidine. This work highlights the potential of spermidine as a vaccine adjuvant in the older adults.

## Results

First we optimised a flow cytometry-based assay to reliably and reproducibly measure autophagy, before applying it to measure autophagy after *in vivo* antigen stimulated T cells post-vaccination. To inhibit the autophagic flux and thereby degradation of LC3-II, the lysosomal inhibitor bafilomycin A was added to the culture for 2 hrs before washing out non-membrane bound LC3-I and staining. In the human lymphocyte line Jurkat cells we confirmed that treatment with bafilomycin A1 increases LC3-II by both flow cytometry and Western blot (Supplementary Fig. S1a and b). Similar results were obtained in PBMC stimulated in vitro (Supplementary Fig. S1c and d). Next we used *in vitro* stimulated control PBMC obtained from five young donors (22-50 yrs old), bled on three different occasions, 2 weeks apart, were stimulated with either IFNγ/ LPS, anti CD40/ IgM or antiCD3/CD28 to induce autophagy in monocytes (Supplementary Fig. S2a), B cells (Supplementary Fig. S2b), CD4+ T (Supplementary Fig. S2c) or CD8+ T (Supplementary Fig. S2d) cells, respectively.

**Figure S1.**
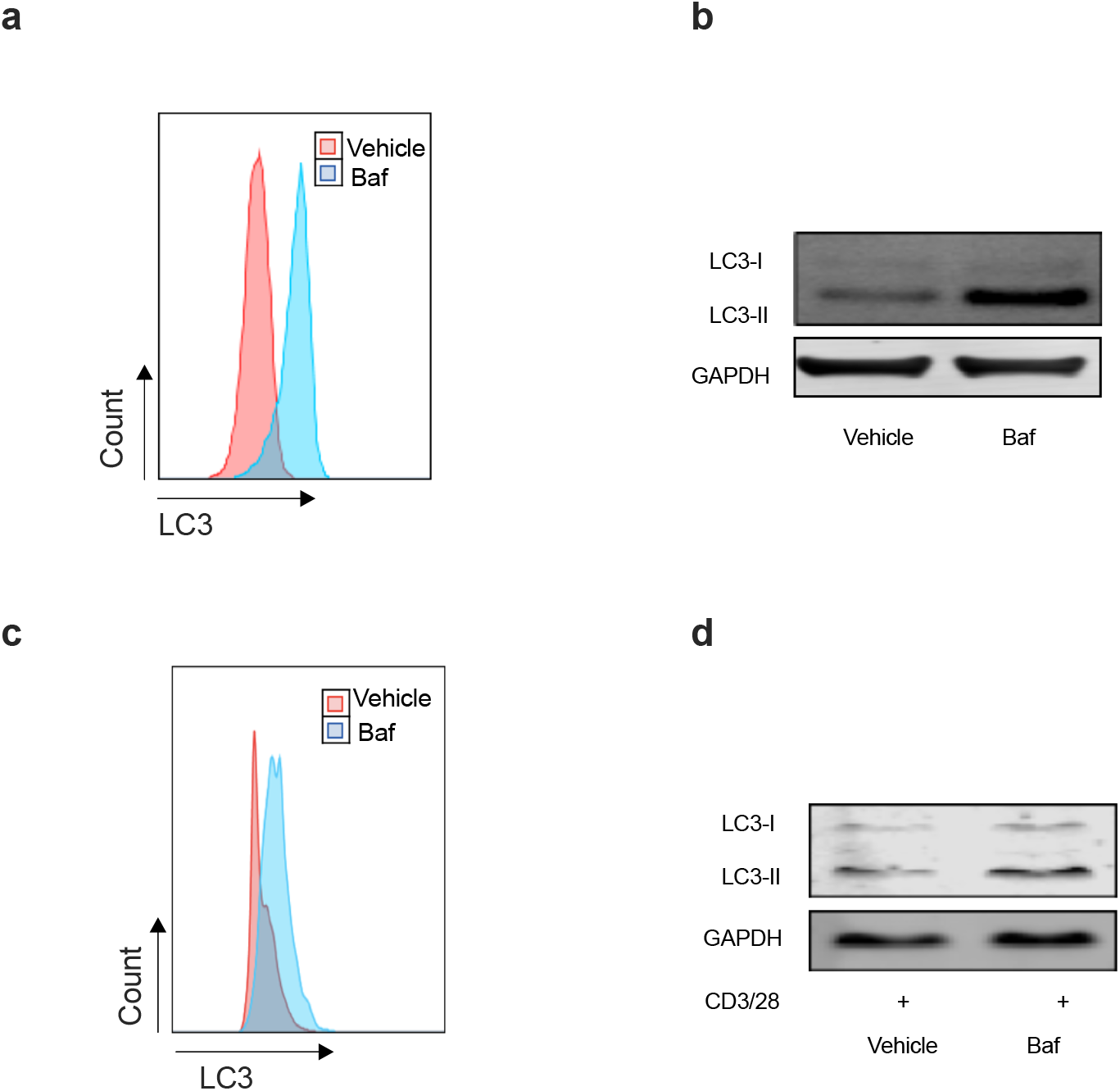
Autophagy levels by flow cytometry-based assay and conventional LC3 western blot in Jurkat cell line and PBMC. **(a-b)** Human T cell line Jurkat was cultured for 24h and treated with or without Bafilomycin A1 for the last 2h. Cells were split into two aliquots, (a) representative flow cytometry-based assay (b) representative western blot for LC3-II and GAPDH for the same sample. (c-d) PBMC from young human donors were cultured with anti-CD3/CD28 for 3 days in the absence/presence of Bafilomycin A1 for the last 2h. Cells were split into two aliquots, (c) representative flow cytometry-based assay (d) representative western blot for LC3-II and GAPDH for the same sample.

**Figure S2.**
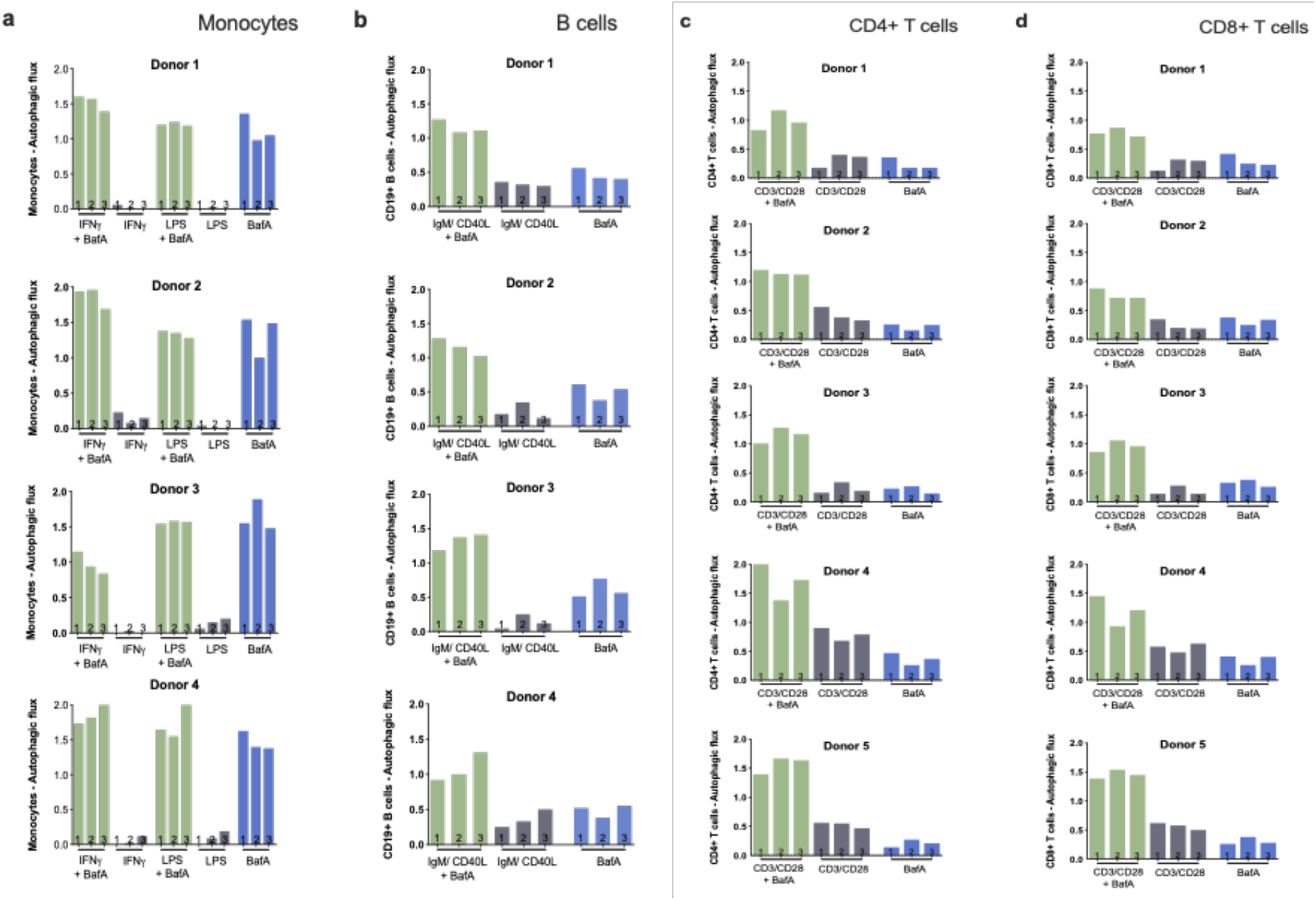
LC3-II detection by flow cytometry is a reliable and reproducible technique in immune cells over several blood draws. PBMC were generated from blood taken at 3 weeks intervals (samples 1, 2, 3) from young human donors and were cultured for 24 hours, in the abence/presence of Bafilomycin A1 for the last 2h. Here basal autophagic flux was calculated as LC3-II mean fluorescence intensity (treatment-basal)/basal. (**a**) Monocytes gated on CD14+ treated with IFNg or LPS, (**b**) B cells gated on CD19+ treated with anti-CD40L and anti-IgM, (**c**) CD4+ T cells gated on CD3+CD4+ treated with anti-CD3/CD28, (**d**) CD8+ T cells gated on CD3+ CD8+ treated with anti-CD3/CD28.

Taken together, the data show: a) autophagic flux varies little within one individual between blood draws; b) there is limited variation between individuals; c) bafilomycin A treatment leads to accumulation of LC3-II in all cell types; d) the respective stimulations via TCR/BCR/TLR or IFNγ-R weakly induce autophagic flux which is consistently further increased in the presence of bafilomycin A.

Published human studies so far have limited their analysis of TCR-induced autophagy to non-specific stimulation *in vitro* such as anti-CD3/CD28. Here we took advantage of an existing cohort of healthy human donors that were vaccinated with a candidate HCV vaccine encoding HCV non-structural proteins (NSmut). Healthy volunteers received an adenoviral vector prime vaccination (ChAd3-NSmut or Ad6-NSmut) followed by a heterologous adenoviral boost vaccination (ChAd3-NSmut or Ad6-NSmut) or an MVA-NSmut boost vaccination (Fig 1a and Supplementary Fig S3a) ^16,17^.

**Figure 1.**
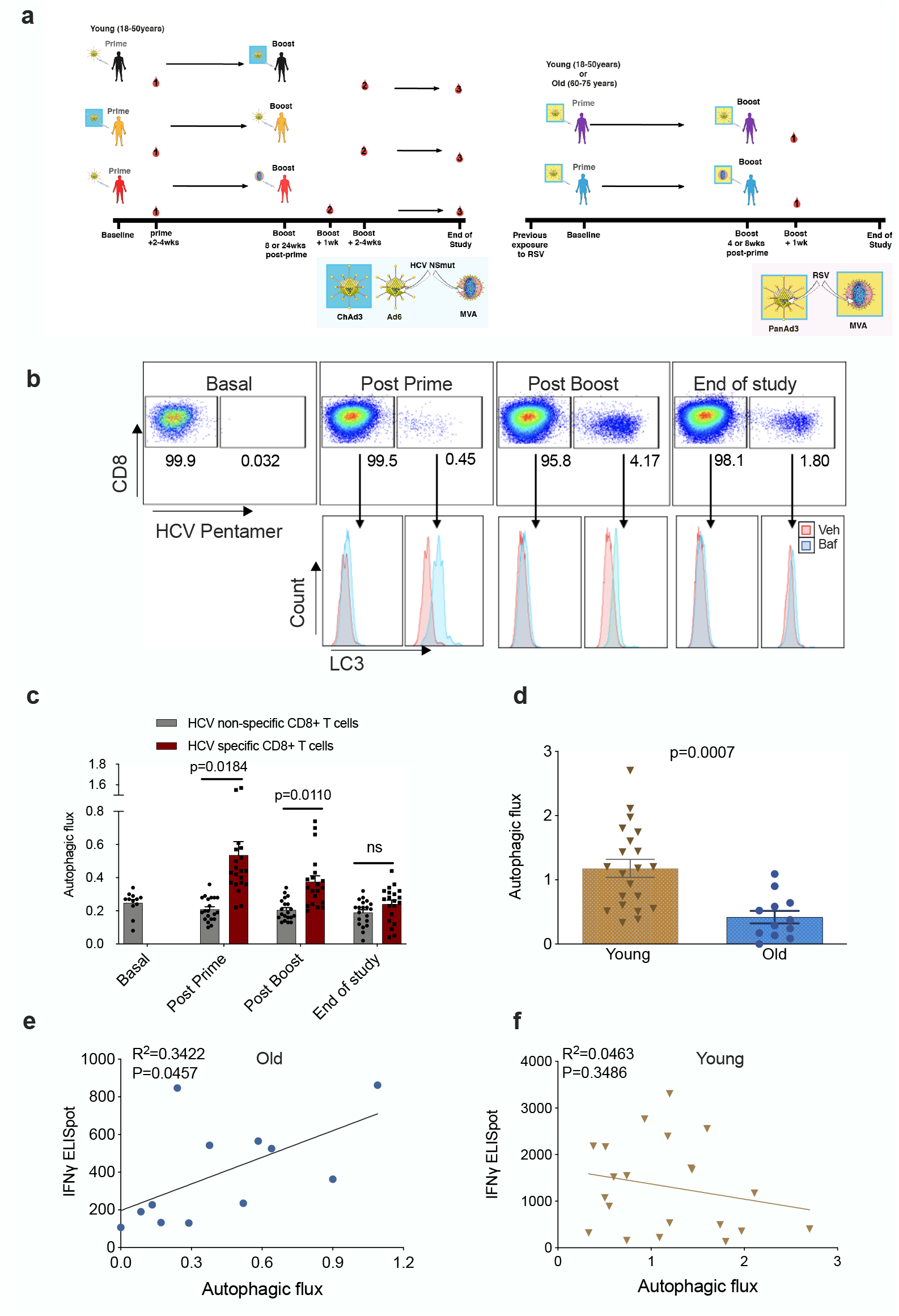
Autophagy is induced by vaccination in antigen-specific T cells and correlates with donor age. Peripheral blood mononuclear cells were isolated from blood samples of vaccinated healthy donors. LC3-II was measured in CD8+ cells using flow cytometry after 2 h treatment with 10 nM bafilomycin A1 (BafA1) or vehicle. Autophagic flux was calculated as LC3-II mean fluorescence intensity (BafA1-Vehicle)/Vehicle. (**a)** Vaccine regimen for HCV and RSV trials (**b**) Representative plots showing Baf A in light blue and vehicle pink (**c**) Quantification in HCV non-specific CD8+ T cells and HCV specific CD8+ T cells detected by HCV pentamers from 10 vaccinees (includling duplicates) using different HCV vaccine regimens, priming with ChimAd and boosting with MVA or AD6 vectors. Autophagy was measured at the peak of the T cell response post prime vaccination, peak of the T cell response post boost vaccination and at the end of the study. (**d**) Autophagic flux was measured in CD8^+^ cells from young donors (N=12, <65yrs) and old donors (N=21, >65 years) vaccinated with respiratory syncytial virus (RSV) 7 days after last boost, quantification calculated as mentioned above. Data represented as mean ± SEM. (**e, f**) Correlation of autophagic flux with total response to peptide pools specific T cell IFNg response to RSV exposure measured by ELISpot in CD8^+^ cells from old donors (**e)** and young donors **(f**), donors as in (**d**). Linear regression with 95% confidence intervals from old and young donors. The goodness of fit was assessed by R^2^. The P value of the slope is calculated by F test.

**Figure S3.**
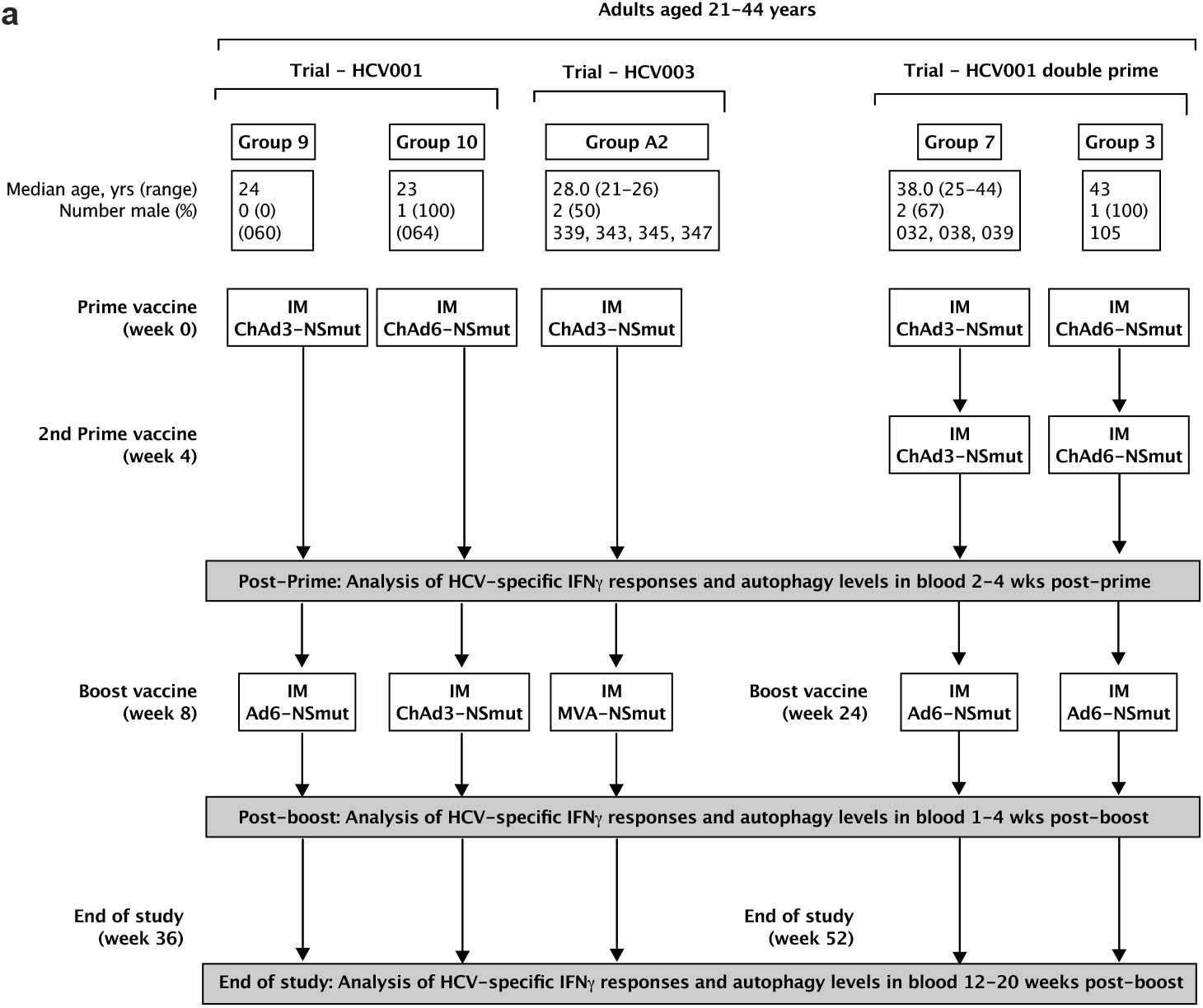
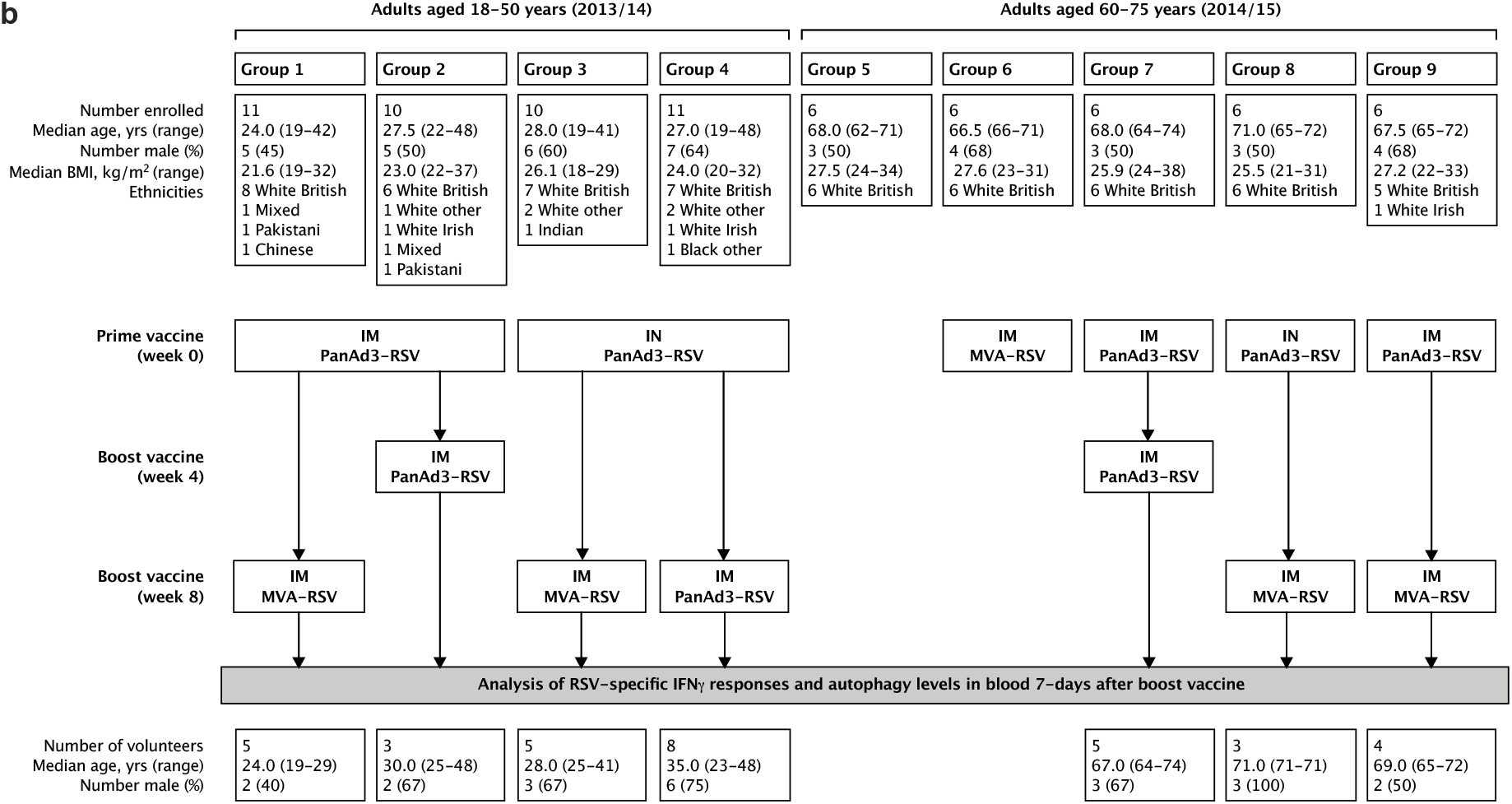
Regimen of immunisations and blood sampling. (**a**) HCV= Hepatitis C virus, ChAd=Chimpanzee Adenoviral Vector, MVA= Modified Ankara Virus vector, (**b**) RSV=respiratory syncitial virus, ChAd=Chimpanzee Adenoviral Vector, MVA= Modified Ankara Virus vector, Unlike for HCV, the adults in the RSV study will have prior immune responses that have been boosted by natural exposure throughout life. In the context of RSV, we still use the term “prime” to mean the first dose of vaccine. Similarly, the term “boost” means the second dose of vaccine and not exposure.

To identify HCV specific CD8+ T cells, PBMC were co-stained with an MHC class I pentamer (HLA-A*02:01, HCV peptide KLSGLGINAV) and anti-LC3-II (Fig 1b). When measured at the peak magnitude of the T cell response to vaccination (2-4 weeks post Ad, 1 week post MVA) HCV specific CD8+ T cells show a significant increase in autophagic flux that is not observed in HCV nonspecific CD8+ T cells (Fig 1c). In antigen-specific T cells, autophagic flux is highest shortly after vaccination but had declined to levels equivalent to antigen-non-specific T cells cells by the end of the study (week 36 or 52). Together, these data show that antigen exposure induces autophagic flux in CD8+ T cells in humans *in vivo*. In previous work we found autophagic flux was reduced in CD8+ T cells from 24 month old mice ^12^. To test whether this is true in human CD8+ T cells from human individuals, and whether this correlates with vaccine immunogenicity, we measured autophagic flux in vaccinees of various ages that were given an experimental RSV vaccine (Fig1a and Supplementary Fig S3b).

As older adults are particularly susceptible to severe disease from RSV infection, the vaccine was given to two age groups (18-50 years and 60-75 years of age). As expected, the naturally-derived population of RSV-specific IFNγ producing CD4+ and CD8+ T cells in peripheral circulation in response to the infection declines with age (Fig. S4).

**Figure S4.**
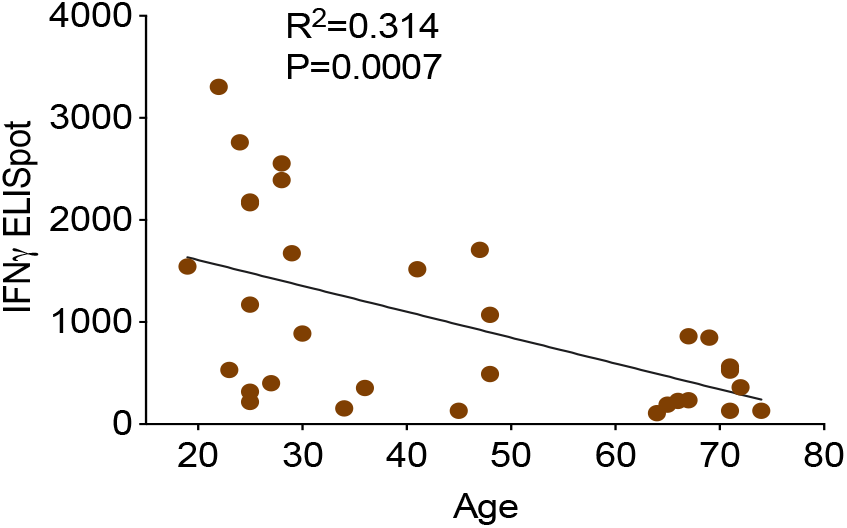
Correlation of age with total and peptide-pool specific T cell IFNγ response to RSV exposure measured by ELISpot in CD8^+^ cells, donors as in Fig 1E. (linear regression with 95% confidence intervals from old and young donors. The goodness of fit was assessed by R^2^. The P value of the slope is calculated by F test.

No MHC class I pentamers were available for RSV to identify antigen-specific T cells, however, bulk T cells of the >60year vaccinees showed significantly lower basal autophagic flux (Fig. 1d). We correlated autophagy levels in T cells with IFNγ ELISpot responses in individual vaccinees and found a strong inverse correlation in the aged group (Fig. 1e) between autophagy and IFNγ responses, but not in the young group (Fig. 1f). Taken together these data suggest that reduced T cell autophagy in aged T cells may underpin reduced T cell responses to vaccination.

We have recently shown that treating old but not young mice with the metabolite spermidine improves autophagy levels in B lymphocytes due to an age-related decline of endogenous spermidine ^15^. Here we sought to confirm this in human lymphocytes. Firstly, we determined spermidine and putrescine levels in PBMC by gas chromatography-mass spectrometry (GC-MS), and found an inverse correlation between age and spermidine but not with putrescine (Fig. 2a). We hypothesized that low levels of spermidine are responsible for low levels of autophagy and poor T cell function in PBMC from old donors. We therefore tested whether supplementation with spermidine recovers T cell autophagy and function. As activation of PBMC with anti-CD3/CD28 optimally induces autophagy levels on day 4 ^18^, we activated PBMC from old donors in the presence of spermidine for 4 days and tested their autophagic flux and function by flow cytometry. Both autophagic flux and the secretion of IFNγ measured by ELISA was improved significantly in T cells from older vaccinees (Fig. 2b and c), Similarly, increased IFNγ can be detected after spermidine treatment by intracellular staining for flow cytometry (Fig. 2d). Interestingly, spermidine supplementation also increases the expression of Perforin (Fig. 2e) but not of Granzyme B (Fig. 2f).

**Figure 2.**
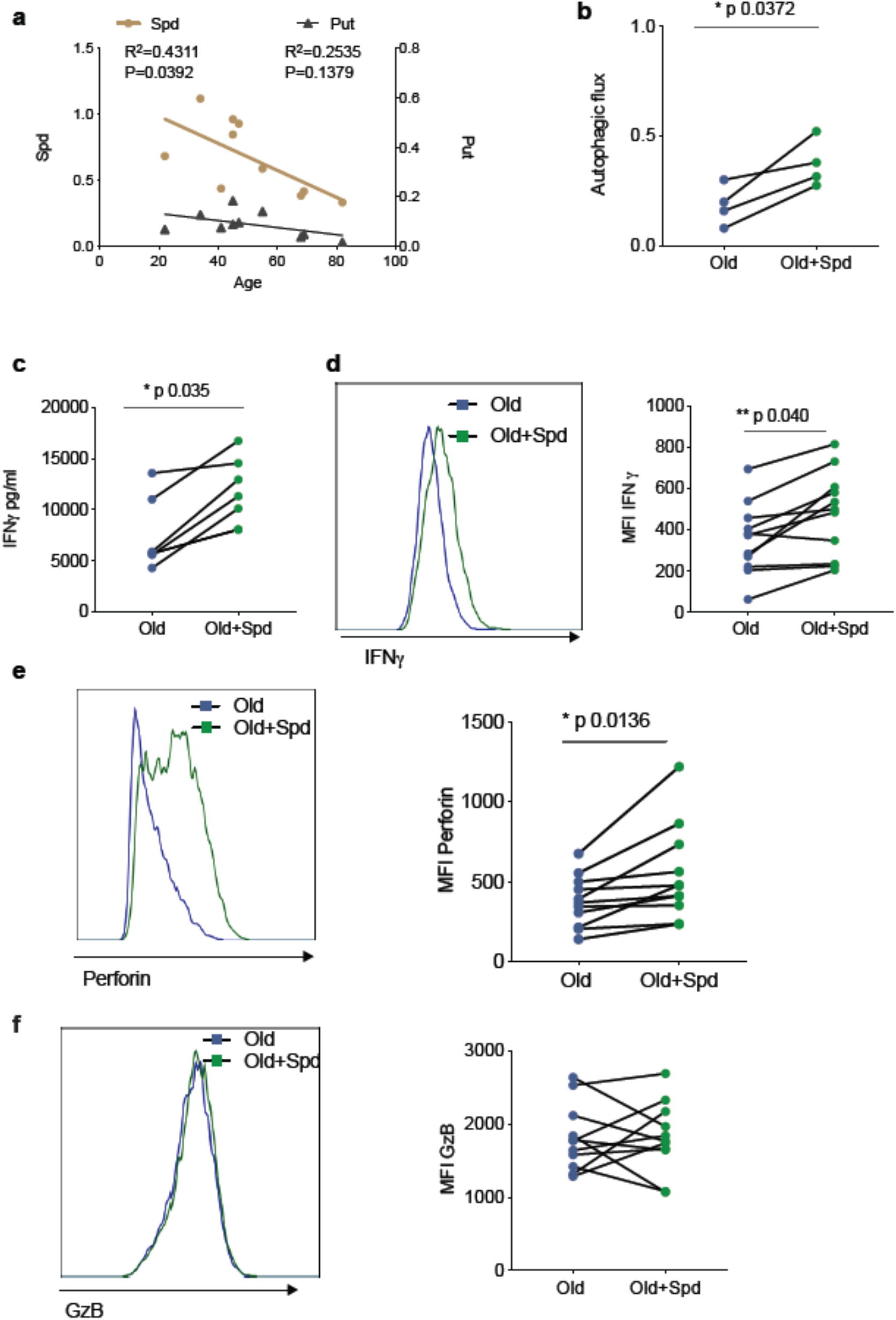
Spermidine declines with age, and supplementing spermidine improves autophagy and CD8+ T cell function in old donors. (**a**) Spermidine and Putrescine content of PBMC collected from healthy donors was measured by GC-MS. Linear regression with 95% confidence intervals. The goodness of fit was assessed by R^2^. The p value of the slope is calculated by F test. (**b-f**) Sorted PBMCs from old human donors (>65yrs) were cultured with anti-CD3/CD28 for 4 days and where indicated treated with 10 μM spermidine, and autophagic flux measured by flow cytometry (**b**), IFNγ assessed by ELISA in tissue culture supernatant (**c**), intracellular IFNγ by flow cytometry (**d**), intracellular Perforin by flow cytometry (**e**), Intracellular Granzyme B **(f)**, all gated on CD8+ cells. Data represented as mean ± SEM, MFI=mean fluorescence intensity. Statistics by paired t-test for b,c,d, e, f.

Next, we investigated whether endogenous polyamine maintains autophagy levels in activated CD8+ T cells from young donors. The drug DFMO (α-difluoromethylornithine) inhibits polyamine biosynthesis by irreversibly inhibiting ornithine decarboxylase (ODC). DFMO almost completely blocked autophagy in CD8+ T cells activated over 7 days, however, when cells were supplemented with spermidine autophagic flux was recovered (Fig. 3a). DFMO also partially blocks IFNγ and Perforin expression in anti-CD3/CD28 activated CD8+ T cells, which can also be rescued by spermidine (Fig. 3b and c). As before, Granzyme B was not affected by DFMO + spermidine (Fig 3d). T cells from young donors, with their high endogenous spermidine levels, do not respond to spermidine supplementation (Fig. 3e, f and g).

**Figure 3.**
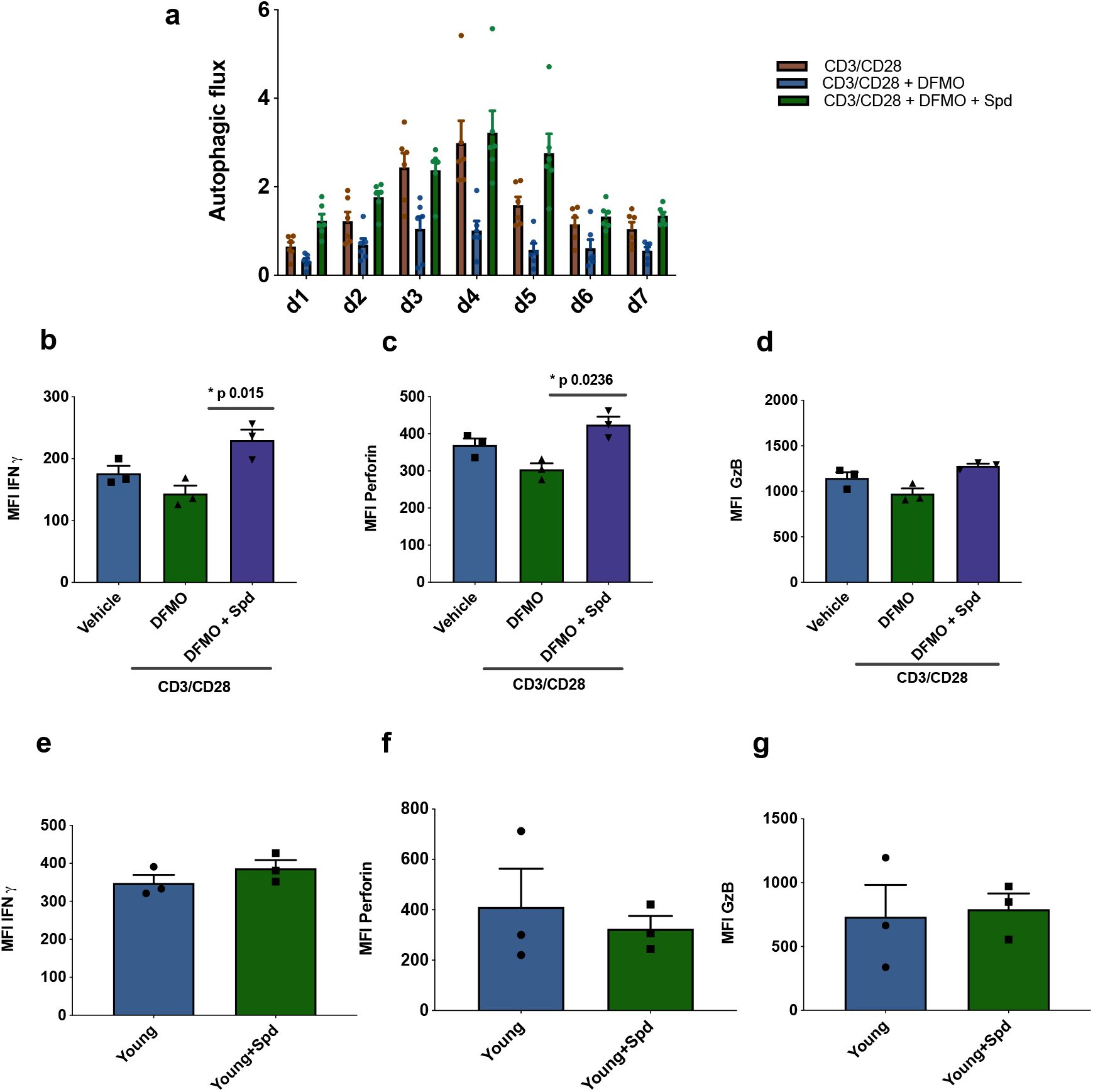
Endogenous spermidine maintains levels of autophagy and T cell function. (**a-d**) Sorted PBMC cells from young human donors (< 65yrs) were activated with anti-CD3/CD28 for 7 days and treated with spermidine synthesis inhibitor DFMO alone or together with 10 μM spermidine. Autophagic flux (**a**) was assessed each day and IFNγ (**b**), Perforin (**c**), Granzyme B (**d**) were measured by flow cytometry in CD8+ cells on day 4. (**e-f**) Sorted PBMC cells from young human donors (< 65yrs) were cultured with anti-CD3/CD28 for 4 days and treated with 10 μM spermidine. (**e**) intracellular IFNγ, (**f**) intracellular Perforin, (**g**) and intracellular Granzyme B were measured in CD8+ cells by flow cytometry. Data represented as mean ± SEM.

We previously found that spermidine maintains autophagic flux via hypusination of eIF5A and TFEB ^15^, and we sought to test this pathway in human T cells. Spermidine lends a moiety to the translation factor eIF5A, which in addition to its role in initiation and termination also promotes the translation of polyproline rich domains, which are difficult to translate ^19^. One such triproline motif-containing protein is TFEB, with mouse TFEB containing one triproline motif while human TFEB contains two. TFEB is the key master transcription factor of autophagosomal and lysosomal gene expression ^20-22^. Here we addressed whether this pathway operates in human T cells and accounts for the loss of autophagy and T cell function. We first verified in the human lymphocyte line Jurkat that the inhibitor GC7 inhibits the hypusinated/ activated form of eIF5A (Fig. 4a) and also confirmed that it decreases LC3-II expression in a dose-dependent manner (Fig. 4b). GC7 reduces TFEB and hypusinated eIF5A in CD8+ T cells activated for 4 days (Fig 4c). It also diminishes the autophagic flux in activated CD8+ T cells over a time course of 7 days (Fig. 4d). We then tested whether spermidine maintains eIF5A hypusination and TFEB levels in CD8+ T cells from young donors by depleting endogenous spermidine with DFMO. While anti-CD3/CD28 increases expression levels of TFEB and eIF5A, a reduction of hypusinated eIF5A and TFEB levels in PBMCs treated with DFMO was observed, which is rescued with spermidine (Fig. 4e). As expected, spermidine treatment of PBMC from young donors demonstrated to have high levels of spermidine does not induce this pathway (Supplementary Fig. S5a) nor T cell function (Fig. 3e, f and g). Finally, we sought to address whether naturally low endogenous spermidine levels can be rescued in PBMC from old donors activated with anti-CD3/CD28. Again, we observed that activation induces levels of protein expression of both hypusinated eIF5A and TFEB, and spermidine further improves both eIF5A and TFEB two-fold (Fig 4f).

**Figure 4.**
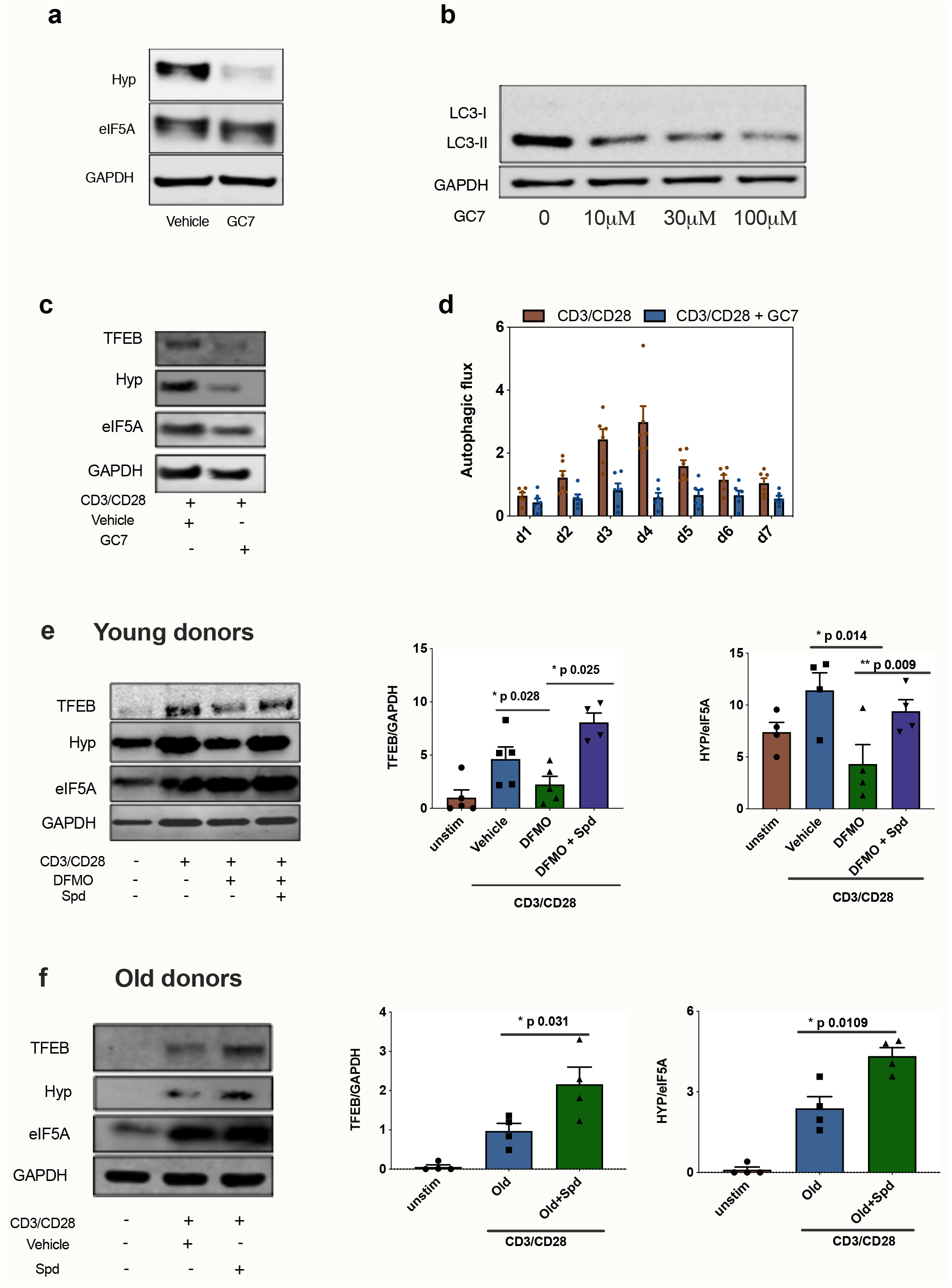
Spermidine’s mode of action is via eIF5A and TFEB in human CD8+ T cells. (**a**) Human T cell line Jurkat was cultured for 24h with 100 μM GC7, then eIF5A and hypusinated eIF5A were measured by WB. (**b**) Jurkat cell line was stimulated with increasing concentrations of GC7 and cell lysates blotted for LC3B. (**C, D**) PBMC from young human donors were cultured with anti-CD3/CD28 for 7 days and treated with GC7. The protein levels of TFEB and eIF5A hypusination were measured in CD8+ cells by Western blot on day 4 (**c**) and autophagic flux was determined as in Fig 1 (**d**). (**e**) PBMC from young human donors were cultured with anti-CD3/CD28 for 4 days and treated with spermidine synthesis inhibitor DFMO alone or together with 10 μM spermidine. The protein levels of TFEB and eIF5A hypusination were measured in CD8+ cells by Western blot. (**f**) Sorted PBMCs from old human donors (>65yrs) were cultured with anti-CD3/CD28 for 4 days and where indicated treated with 10 μM spermidine, The protein levels of TFEB and eIF5A hypusination were measured in CD8+ cells by Western blot. Target band intensity was normalized to eIF5A (for Hyp) or GAPDH (for TFEB). Data represented as mean ± SEM.

**Figure S5.**
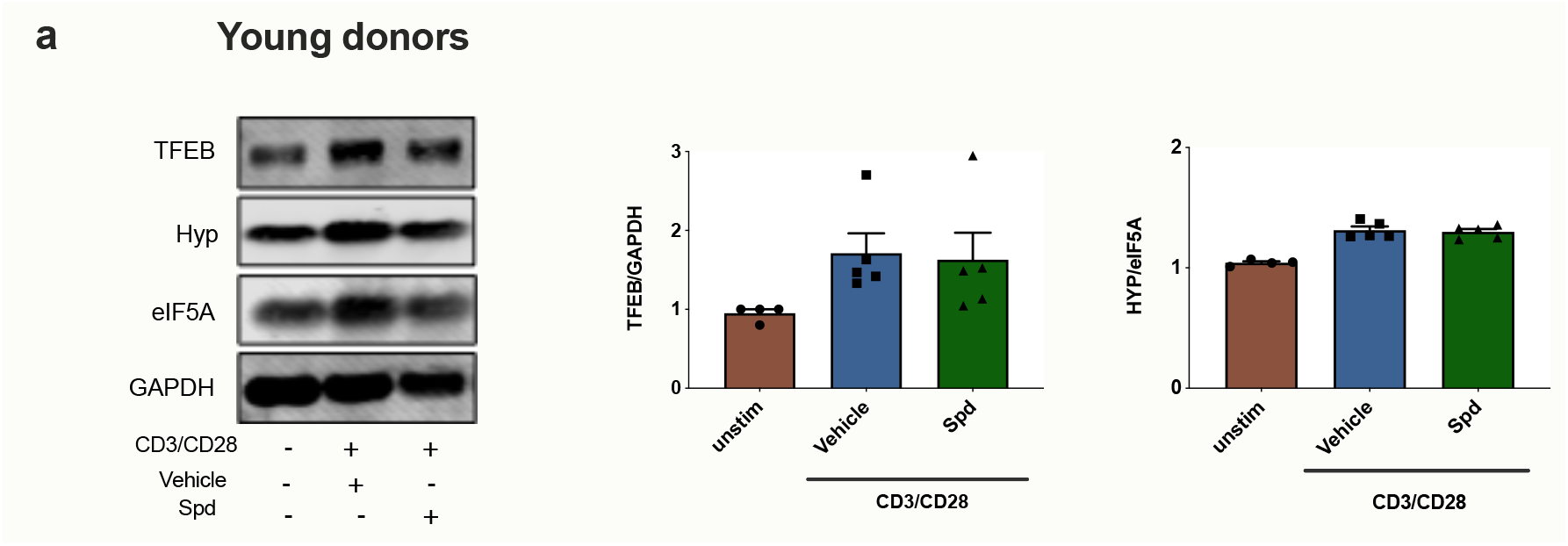
Spermidine does not improve eIF5A and TFEB in young donors. (**a**) PBMC from young human donors were cultured with anti-CD3/CD28 for 4 days and treated with 10μM spermidine for 4 days. The protein levels of TFEB and eIF5A hypusination were measured in sorted CD8+ cells by Western blot. Target band intensity was normalized to eIF5A (for Hyp) or GAPDH (for TFEB). Data represented as mean ± SEM. *

## Discussion

Several studies have addressed the induction of autophagy by antigenic stimulation in murine T cells in vivo and in response to anti-CD3/CD28 in vitro stimulation of human T cells ^23^. In mouse studies Xu et al measured a peak in autophagy induction shortly after the CD8+ effector phase with subsequent failure to mount memory responses ^11^. In line with this, we and Xu et al showed that deletion of autophagy mostly affects T cells in the memory phase of CD8+ T cell responses as opposed to the effector phase ^11,12^. The question remains whether it is initiation or maintenance of memory responses that are affected. Interestingly, the work of Schlie et al showed that the memory response could be rescued by N-acetyl cysteine, arguing that memory cells may be formed but not maintained without autophagy ^24^. While it has been shown in human T cells that autophagy peaks after 4 days of anti-CD3/CD28 stimulation, later time points are more difficult to mimic accurately in vitro. Here we show for the first time that antigenic encounter in vivo induces autophagy after the effector phase, arguing that autophagy may be important in human CD8+ T cells during the memory phase. Interestingly, we could not observe any bystander effect on CD8+ T cells that are not specific for HCV, suggesting that it is a cell-intrinsic effect through specific stimulation via their TCR.

The role of autophagy in these long term surviving antigen-specific T cells is still elusive: autophagy could either degrade molecules no longer needed such as PU1 in the establishment of a Th9 response ^25^ or CDKN1 in the effector phase of CD8 T cells ^26^ or organelles such as mitochondria or ER ^12,27^. Alternatively, autophagy could provide building blocks including amino acids, free fatty acids and nucleotides that are important for essential cellular requirements: provision of ATP ^28,29^, transcription and translation for T cell function. Or a combination of these mechanisms may be at play here, something which remains to be elucidated.

With increasing life expectancy, the number of people over 60 years of age is expected to double by 2050, reaching 2.1 billion worldwide. The severity of many infections is higher in the older population compared to younger adults as particularly notable during the SARS2-CoV pandemic. Moreover, the success of childhood vaccination is widely recognized but the importance of vaccination of the older population is frequently underestimated ^3^. Immune responses to vaccines are known to be particularly ineffective in the older population and yet some vaccines such as for influenza and SARS2-Cov are primarily needed for the older adults. The development of drugs that improve vaccination in this expanding population is therefore an urgent socio-economic need.

Targeting mTOR with Rapamycin was the first drug both in mice and in human clinical trials shown to have a beneficial effect on T cell responses in mice and older humans ^30,31^. However, whether Rapamycin triggers autophagy at the administered dose has not been investigated. With this study we show for the first time that the autophagy-inducing drug spermidine has an immune boosting effect on the T cell compartment in humans in vitro and that low autophagy levels correlate with low responses to vaccination. The translation of TFEB is one of the limiting factors for sufficient autophagy levels required to mount an immune response. In addition, to be active, TFEB needs to be dephosphorylated for its translocation into the nucleus, which mTOR inhibition facilitates ^32-34^. Therefore, spermidine and an mTOR inhibitor may have to be combined to optimally restore immune responses in the older adults.

We find that spermidine levels decline in PBMCs in the older humans. This confirms earlier study in plasma in which spermidine levels were found to be low in the >65 age group and rising again in centenarians ^35^. Studies of this kind usually indicate that the phenotype is maladaptive with age as centenarians do not display many of the aging features of the age group below. However, the reasons for the age-related decline in spermidine in blood cells are not clear, and currently under investigation. Overall it is evident that endogenous spermidine maintains autophagy in human T cells, a novel metabolic pathway that has not been investigated before.

Spermidine has recently been administered to humans in a small experimental trial with a beneficial effect on cognitive function without adverse effects ^36,37^. It remains to be shown whether at such low doses it has an effect on autophagy. Our study strongly suggests that a small experimental trial should be conducted to test whether spermidine can be used to improve vaccination efficiency in older adults - eIF5A, TFEB and autophagy in PBMCs could be used as biomarkers to determine dose, duration and biomarkers for the anti-immune senescence effect.

In conclusion we validated a novel anti-immune senescence pathway in humans with druggable targets and biomarkers, which could also be used for other broader antiaging drug trials.

## Materials and Methods

**Human Samples** Human peripheral blood mononuclear cells (PBMC) were obtained under the ethics reference NRES Berkshire 13/SC/0023, from phase I clinical trials of novel viral-vectored vaccines for hepatitis-C virus (HCV;

NCT01070407 and NCT01296451) or respiratory syncytial virus (RSV), described in more detail elsewhere ^16,17 38 39 40^. Volunteers were self-selected adults who provided written informed consent and who were carefully screened for being healthy before vaccination. The vaccine schedules are described in Diagrams (Fig S4 A and B). Blood samples were collected in heparinised tubes for assays that required PBMC. PBMC were isolated within 6 h of sample collection. An aliquot of PBMC was immediately used for fresh ELISpot assays and the remainder cryopreserved in Recovery^™^ Cell Freezing Medium. Serum samples were obtained by centrifugation of whole blood collected in clotted tubes, and then cryopreserved.

**Control PBMC** were isolated from blood or blood cones of healthy donors using Ficoll-Paque density gradient separation. All volunteers provided written informed consent. The study was approved by the Local Ethics Committee Oxford and Birmingham. Freshly isolated PBMCs were cultured directly or were frozen in 90% FBS and 10% DMSO in liquid nitrogen. Fresh or thawed PBMCs were cultured with RPMI 1640, 10% FCS, 2 mM L-Glutamine, 100 U/mL Penicillin (Invitrogen).

### ELISpot Assay

*Ex vivo* IFNγ ELISpot assays were performed according to manufacturers’ instructions (Mabtech) on freshly isolated PBMC plated in triplicates at 2×10^5^ PBMC per well. In brief, peptide pools consisted of mainly 15-mer sequences with 11 amino acid overlaps and covering the sequence of proteins F, N and M2-1. Peptides were dissolved in 100% DMSO and arranged in four pools. DMSO (the peptide diluent) and Concanavalin A (ConA) were used as negative and positive controls, respectively. The mean + 4 StDev of the DMSO response from all samples identified a cut off whereby individual samples with background DMSO values ≥50 spot forming cells per million PBMCs were excluded from analysis. Calculation of triplicate well variance was performed as described previously^40^.

### Human T cell assays

The mean age of young donors was 40.7±11.3 years and the mean age of old donors was 77.6±6.6. PBMC were activated with either soluble anti-CD3 (1 μg/ml, Jackson Immuno Research) and anti-CD28 (1 μg/ml, Jackson Immuno Research) with or without 10μM spermidine, or 1 mM difluoromethylornithine (DFMO, Enzo Life Sciences) or 10 μM GC7 (or as indicated, Millipore) for 4 days. After MACS sorting of CD8+ T cell using a negative selection kit (CD8+ T Cell Isolation Kit II, human, Miltenyi Biotec), cells were lysed for western blotting or stained for Autophagy flux assay as described below. IFNγ release in culture supernatants was measured by heterologous two-site sandwich ELISA, according to the manufacturer’s protocol (Invitrogen).

### Autophagy Flux Assay for Flow Cytometry

PBMC from healthy donors, activate for T cells with CD3/CD28 beads (Dynabeads Thermo Fisher) (1:1), for B cells with anti-IgM (5 μg/ml, Jackson Immuno Research) and CD40L (100 ng/ml, Enzo Life science) and for monocytes activated with IFN γ (20 ng/ml, Jackson Immuno Research) and/or LPS (10 μg/ml, Santa Cruz) (all 24hrs except for T cells which were stimulated overnight).

Autophagy levels were measured after 2h treatment with bafilomycin A1 (10nM BafA1) or vehicle. We adapted the FlowCellect Autophagy LC3 antibody-based assay kit (FCCH100171, Millipore) as follows: In brief, cells were stained with surface markers (as above) and washed with Assay Buffer in 96 well U bottom plates. 0.05% Saponin was added to each well and spun immediately, followed by anti-LC3 (FITC) at 1:20 in Assay Buffer, (30-50μl/ well) at 4°C for 30 minutes, and washed once with 150μl Assay Buffer. Stained cells were fixed with 2% PFA before FACS analysis. Autophagic flux was calculated as LC3-II mean fluorescence intensity of (BafA1-Vehicle)/Vehicle.

### Surface Staining for Flow Cytometry

For CellTrace staining, CellTrace Violet (C34557, Thermo Fisher) was used according to the manufacturer’s protocol. Cells were transferred to a round bottom 96 well plate and centrifuged (300xg, 5 minutes). The pellet was resuspended in PBS containing the viability dye Live/Dead (Life Technologies) or fixable Zombie Aqua Live/Dead (423102, Biolegend) for 10 minutes in the dark at room temperature (RT) to exclude dead cells during analysis. After washing with PBS/5% FCS, cells were resuspended in PBS/2% FCS/ 5mM EDTA (FACS buffer) containing a cocktail of antibodies relevant to the desired cell surface markers. Fc block was typically added to the antibody mix to minimise non-specific staining. Surface antibody staining was performed at 4°C for 20 mins in the dark. A list of all surface antibodies utilised and their working concentrations are in Table 2. Following incubation cells were washed with FACS buffer and immediately analysed on a four-laser LSR Fortessa X-20 flow cytometer. Acquired data were analyzed using FlowJo 10.2.

### Intracellular Staining for Flow Cytometry

For intracellular staining, PBMC were stimulated in R10 with anti-CD3 (1 μg/ml, Jackson Immuno Research) and anti-CD28 (1 μg/ml, Jackson Immuno Research) with or without 10μM spermidine for 4 days. On day 4 cells were re-stimulated with the same concentrations of anti-CD3/CD28 for 6 hours at 37°C in the presence of 1 μg/ml brefeldin-A (Sigma-Aldrich). As a control, cells were left unstimulated. Following surface marker staining as described above, cells were fixed with 100μL Fixation buffer (eBioscience) for 20 min at RT in the dark. Next, cells were permeabilised with 100μL of 1x Permeabilisation buffer (eBioscience) for 15 min in the dark at RT. Then, cells were resuspended in the intracellular antibody mix (anti-IFN-γ, anti-Granzyme B, anti-Perforin) and incubated for 30 min in the dark at RT. After being washed twice with Permeabilization buffer the cells were resuspended in 200μL of FACS buffer for analysis.

### MHC class I pentamer staining to identify antigen-specific T cells

An HCV-specific HLA-A*02-restricted pentamer, peptide sequence KLSGLGINAV (Proimmune) was used to identify HCV-specific CD8+ T cells *ex vivo*. PBMC were washed in PBS and were stained with pentamers at room temperature (20mins) in PBS, washed twice in PBS before further mAb staining as described above. During analysis, stringent gating criteria were applied with doublet and dead cell exclusion to minimise nonspecific binding contamination.

### Western blot

Cells in suspension were washed with PBS and lysed using NP-40 lysis buffer containing proteinase inhibitors (Sigma) and phosphatase inhibitors (Sigma) on ice. After spinning down the debris, protein concentration in the supernatant was quantified by BCA Assay (23227, Thermo Fisher). Reducing Laemmli Sample Buffer was then added to the supernatant and heated at 100°C for 5 minutes. 5-20 *μ*g protein was loaded for SDS-PAGE analysis. NuPAGE Novex 4-12% Bis-Tris gradient gel (Thermo Fisher) with MES running buffer (Thermo Fisher) was used. Proteins were transferred to a PVDF membrane (IPFL00010, Millipore) and blocked with 5% skimmed milk-TBST. Membranes were incubated with primary antibodies dissolved in 1% milk overnight and secondary antibodies dissolved in 1% milk with 0.01% SDS for imaging using the Odyssey CLx Imaging System. Data were analyzed using Image Studio Lite.

### Spermidine measurement in cell lysates by GC-MS

This protocol was used as published previously ^41^. Briefly, cells were washed with PBS and the pellet resuspended in lysis buffer (80% methanol + 5% Trifluoroacetic acid) spiked with 2.5 μM 1,7-diaminoheptane (Sigma). The cell suspension, together with acid-washed glass beads (G8772, Sigma), was transferred to a bead beater tube and homogenized in a bead beater (Precellys 24, Bertin Technologies) for four cycles (6500 Hz, 45 s) with 1 minute of ice incubation between each cycle. The homogenized samples were centrifuged at 13,000 g for 20 minutes at 4°C. The supernatant was collected and dried overnight. For chemical derivatization, 200 μL trifluoroacetic anhydride was added to the dried pellet and incubated at 60°C for 1 hour, shaking at 1200 rpm. The derivatization product was dried, re-suspended in 30 μL isopropanol and transferred to glass loading-vials. The samples were analyzed using a GCxGC-MS system as described ^41^. The following parameters were used for quantification of the 1D-GC-qMS data: Slope: 1000/min, width: 0.04 s, drift 0/min and T. DBL: 1000 min without any smoothing methods used.

### Statistical analyses

Prism software (GraphPad) was used for statistical analyses. Data are represented as mean ± SEM. All comparative statistics were post-hoc analyses. Paired or unpaired two-tailed Student’s t-test was used for comparisons between two normally distributed data sets with equal variances. Linear regression with a 95% confidence interval was used to assess the relationships between age and the expression of target proteins or spermidine levels, in which R^2^ was used to assess the goodness of fit and the P value calculated from F test was used to assess if the slope was significantly non-zero. P values were used to quantify the statistical significance of the null hypothesis testing. *P≤0.05, **P≤0.01, ***P≤0.001, ****P≤0.0001, ns, not significant.

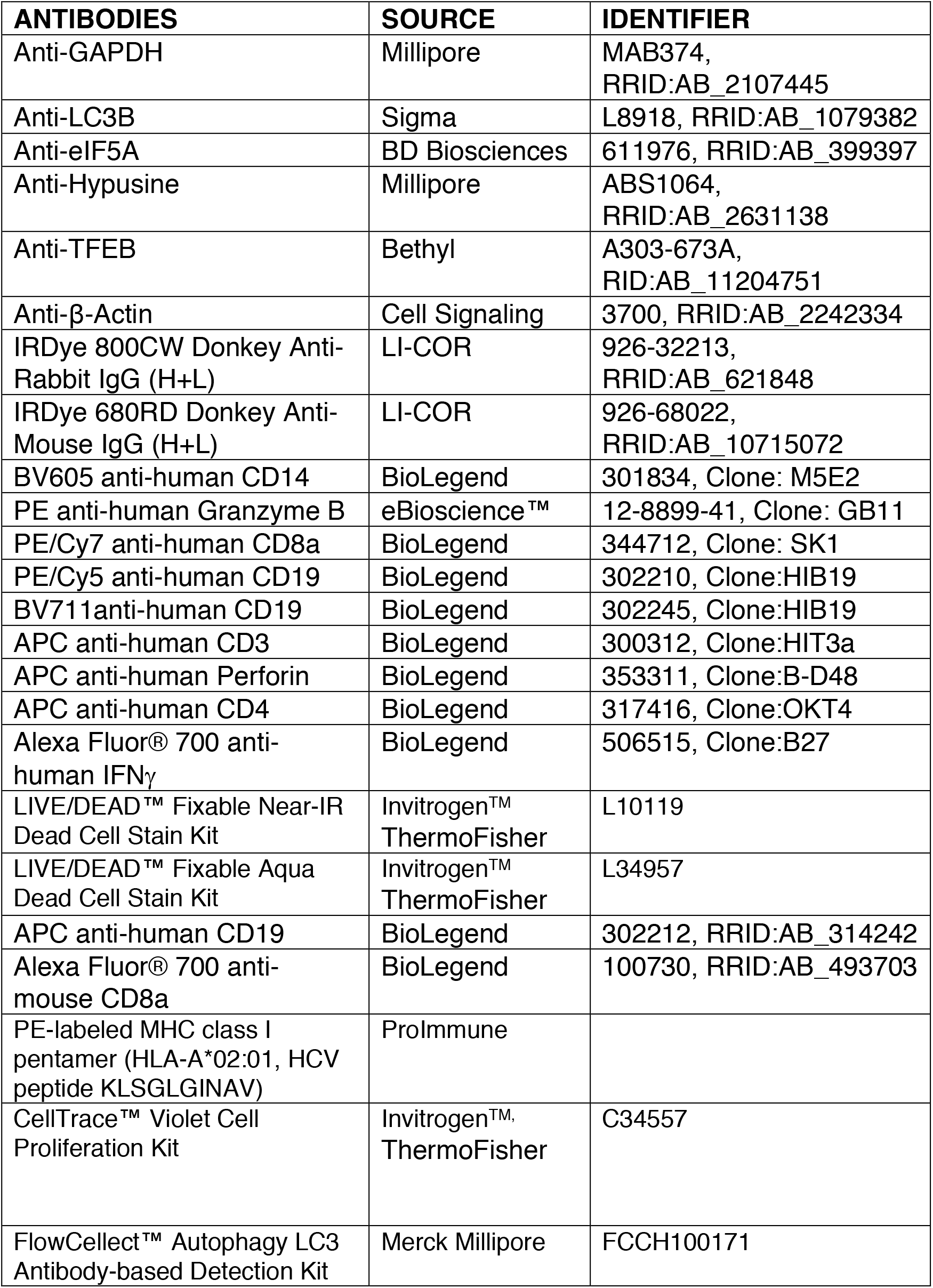

## Acknowledgements

We would like to acknowledge Reithera/Okairos for RSV trial, Zhanru Yu for Spermidine and Putrescine measurements, Kirsty McGee and Christos Ermogenous for ethical approval and provision of samples from Birmingham healthy older donors. We would like to thank all donors, volunteers and phlebotomists. K.S. is a Wellcome Investigator, which funded this study. Wellcome funding (WT109665MA) and NIHR SF was also awarded to P.K. We acknowledge funding from the National Institute for Health Research (NIHR) Oxford Biomedical Research Centre (BRC). The views expressed are those of the authors and not necessarily those of the NHS, the NIHR or the Department of Health.

## Contributions

G.A. designed and performed most experiments prepared all figures and contributed to the writing of the manuscript. I.P. set up the autophagy flux protocol in the primary cells and performed the majority of the experiments for the vaccine samples. L.S. helped to prepare the figures and corrected the manuscript. H.Z. performed some experiments and initiated the eIF5A/TFEB project. C.G., E.B. and P.K. conducted the vaccine trials. P.K. and J.L. corrected the manuscript. C.G. gave advice on data interpretation and corrected the manuscript. Healthy human samples were provided by J.L. and A.M.. A.K.S. supervised this project, designed the experiments, provided funds, and wrote the manuscript.

## Reference

1 Zhang, J. et al. Progress and Prospects on Vaccine Development against SARS-CoV-2. Vaccines (Basel) 8, doi:10.3390/vaccines8020153 (2020).

2 Lurie, N., Saville, M., Hatchett, R. & Halton, J. Developing Covid-19 Vaccines at Pandemic Speed. N Engl J Med, doi:10.1056/NEJMp2005630 (2020).

3 Weinberger, B. Vaccines for the elderly: current use and future challenges. Immun Ageing 15, 3, doi:10.1186/s12979-017-0107-2 (2018).

4 Chen, W. H. et al. Vaccination in the elderly: an immunological perspective. Trends Immunol 30, 351–359, doi:10.1016/j.it.2009.05.002 (2009).

5 Zhang, H., Puleston, D. J. & Simon, A. K. Autophagy and Immune Senescence. Trends Mol Med 22, 671–686, doi:10.1016/j.molmed.2016.06.001 (2016).

6 Rubinsztein, D. C., Marino, G. & Kroemer, G. Autophagy and aging. Cell 146, 682–695, doi:10.1016/j.cell.2011.07.030 (2011).

7 Salminen, A., Kaarniranta, K. & Kauppinen, A. Inflammaging: disturbed interplay between autophagy and inflammasomes. Aging (Albany NY) 4, 166–175, doi:10.18632/aging.100444 (2012).

8 Saitoh, T. et al. Loss of the autophagy protein Atg16L1 enhances endotoxin-induced IL-1beta production. Nature 456, 264–268, doi:10.1038/nature07383 (2008).

9 Stranks, A. J. et al. Autophagy Controls Acquisition of Aging Features in Macrophages. J Innate Immun 7, 375–391, doi:10.1159/000370112 (2015).

10 Phadwal, K. et al. A novel method for autophagy detection in primary cells: impaired levels of macroautophagy in immunosenescent T cells. Autophagy 8, 677–689, doi:10.4161/auto.18935 (2012).

11 Xu, X. et al. Autophagy is essential for effector CD8(+) T cell survival and memory formation. Nat Immunol 15, 1152–1161, doi:10.1038/ni.3025 (2014).

12 Puleston, D. J. et al. Autophagy is a critical regulator of memory CD8(+) T cell formation. Elife 3, doi:10.7554/eLife.03706 (2014).

13 Mortensen, M. et al. The autophagy protein Atg7 is essential for hematopoietic stem cell maintenance. J Exp Med 208, 455–467, doi:10.1084/jem.20101145 (2011).

14 Madeo, F., Eisenberg, T., Pietrocola, F. & Kroemer, G. Spermidine in health and disease. Science 359, doi:10.1126/science.aan2788 (2018).

15 Zhang, H. et al. Polyamines Control eIF5A Hypusination, TFEB Translation, and Autophagy to Reverse B Cell Senescence. Mol Cell 76, 110–125 e119, doi:10.1016/j.molcel.2019.08.005 (2019).

16 Swadling, L. et al. Highly-Immunogenic Virally-Vectored T-cell Vaccines Cannot Overcome Subversion of the T-cell Response by HCV during Chronic Infection. Vaccines (Basel) 4, doi:10.3390/vaccines4030027 (2016).

17 Barnes, E. et al. Novel adenovirus-based vaccines induce broad and sustained T cell responses to HCV in man. Sci Transl Med 4, 115ra111, doi:10.1126/scitranslmed.3003155 (2012).

18 Watanabe, R. et al. Autophagy plays a protective role as an anti-oxidant system in human T cells and represents a novel strategy for induction of T-cell apoptosis. Eur J Immunol 44, 2508–2520, doi:10.1002/eji.201344248 (2014).

19 Gutierrez, E. et al. eIF5A promotes translation of polyproline motifs. Mol Cell 51, 35–45, doi:10.1016/j.molcel.2013.04.021 (2013).

20 Napolitano, G. & Ballabio, A. TFEB at a glance. J Cell Sci 129, 2475–2481, doi:10.1242/jcs.146365 (2016).

21 Lapierre, L. R., Kumsta, C., Sandri, M., Ballabio, A. & Hansen, M. Transcriptional and epigenetic regulation of autophagy in aging. Autophagy 11, 867–880, doi:10.1080/15548627.2015.1034410 (2015).

22 Settembre, C. et al. TFEB links autophagy to lysosomal biogenesis. Science 332, 1429–1433, doi:10.1126/science.1204592 (2011).

23 Macian, F. Autophagy in T Cell Function and Aging. Front Cell Dev Biol 7, 213, doi:10.3389/fcell.2019.00213 (2019).

24 Schlie, K. et al. Survival of effector CD8+ T cells during influenza infection is dependent on autophagy. J Immunol 194, 4277–4286, doi:10.4049/jimmunol.1402571 (2015).

25 Rivera Vargas, T. et al. Selective degradation of PU.1 during autophagy represses the differentiation and antitumour activity of TH9 cells. Nat Commun 8, 559, doi:10.1038/s41467-017-00468-w (2017).

26 Jia, W. et al. Autophagy regulates T lymphocyte proliferation through selective degradation of the cell-cycle inhibitor CDKN1B/p27Kip1. Autophagy 11, 2335–2345, doi:10.1080/15548627.2015.1110666 (2015).

27 Jia, W., Pua, H. H., Li, Q. J. & He, Y. W. Autophagy regulates endoplasmic reticulum homeostasis and calcium mobilization in T lymphocytes. J Immunol 186, 1564–1574, doi:10.4049/jimmunol.1001822 (2011).

28 Shi, L. Z. et al. HIF1alpha-dependent glycolytic pathway orchestrates a metabolic checkpoint for the differentiation of TH17 and Treg cells. J Exp Med 208, 1367–1376, doi:10.1084/jem.20110278 (2011).

29 Pearce, E. L. et al. Enhancing CD8 T-cell memory by modulating fatty acid metabolism. Nature 460, 103–107, doi:10.1038/nature08097 (2009).

30 Araki, K. et al. mTOR regulates memory CD8 T-cell differentiation. Nature 460, 108–112, doi:10.1038/nature08155 (2009).

31 Mannick, J. B. et al. TORC1 inhibition enhances immune function and reduces infections in the elderly. Sci Transl Med 10, doi:10.1126/scitranslmed.aaq1564 (2018).

32 Martina, J. A., Chen, Y., Gucek, M. & Puertollano, R. MTORC1 functions as a transcriptional regulator of autophagy by preventing nuclear transport of TFEB. Autophagy 8, 903–914, doi:10.4161/auto.19653 (2012).

33 oczniak-Ferguson, A. et al. The transcription factor TFEB links mTORC1 signaling to transcriptional control of lysosome homeostasis. Sci Signal 5, ra42, doi:10.1126/scisignal.2002790 (2012).

34 Settembre, C. et al. A lysosome-to-nucleus signalling mechanism senses and regulates the lysosome via mTOR and TFEB. EMBO J 31, 1095–1108, doi:10.1038/emboj.2012.32 (2012).

35 Pucciarelli, S. et al. Spermidine and spermine are enriched in whole blood of nona/centenarians. Rejuvenation Res 15, 590–595, doi:10.1089/rej.2012.1349 (2012).

36 Schwarz, C. et al. Safety and tolerability of spermidine supplementation in mice and older adults with subjective cognitive decline. Aging (Albany NY) 10, 19–33, doi:10.18632/aging.101354 (2018).

37 Wirth, M. et al. Effects of spermidine supplementation on cognition and biomarkers in older adults with subjective cognitive decline (SmartAge)-study protocol for a randomized controlled trial. Alzheimers Res Ther 11, 36, doi:10.1186/s13195-019-0484-1 (2019).

38 Green, C. A. et al. Safety and immunogenicity of novel respiratory syncytial virus (RSV) vaccines based on the RSV viral proteins F, N and M2-1 encoded by simian adenovirus (PanAd3-RSV) and MVA (MVA-RSV); protocol for an open-label, dose-escalation, single-centre, phase 1 clinical trial in healthy adults. BMJ Open 5, e008748, doi:10.1136/bmjopen-2015-008748 (2015).

39 Green, C. A. et al. Chimpanzee adenovirus- and MVA-vectored respiratory syncytial virus vaccine is safe and immunogenic in adults. Sci Transl Med 7, 300ra126, doi:10.1126/scitranslmed.aac5745 (2015).

40 Green, C. A. et al. Novel genetically-modified chimpanzee adenovirus and MVA-vectored respiratory syncytial virus vaccine safely boosts humoral and cellular immunity in healthy older adults. J Infect 78, 382–392, doi:10.1016/j.jinf.2019.02.003 (2019).

41 Yu, Z. et al. Optimizing 2D gas chromatography mass spectrometry for robust tissue, serum and urine metabolite profiling. Talanta 165, 685–691, doi:10.1016/j.talanta.2017.01.003 (2017).

